# A Benchmarking Platform for Assessing Protein Language Models on Function-related Prediction Tasks

**DOI:** 10.1101/2025.04.10.648084

**Authors:** Elif Çevrim, Melih Gökay Yiğit, Erva Ulusoy, Ardan Yılmaz, Tunca Doğan

## Abstract

Proteins play a crucial role in almost all biological processes, serving as the building blocks of life and mediating various cellular functions, from enzymatic reactions to immune responses. Accurate annotation of protein functions is essential for advancing our understanding of biological systems and developing innovative biotechnological applications and therapeutic strategies. To predict protein function, researchers primarily rely on classical homology-based methods, which use evolutionary relationships, and increasingly on machine learning (ML) approaches. Lately, protein language models (PLMs) have gained prominence; these models leverage specialised deep learning architectures to effectively capture intricate relationships between sequence, structure, and function. We recently conducted a comprehensive benchmarking study to evaluate diverse protein representations (i.e., classical approaches and PLMs) and discuss their trade-offs. The current work introduces the Protein Representation Benchmark – PROBE tool, a benchmarking framework designed to evaluate protein representations on function-related prediction tasks. Here, we provide a detailed protocol for running the framework via the GitHub repository and accessing our newly developed user-friendly web service. PROBE encompasses four core tasks: semantic similarity inference, ontology-based function prediction, drug target family classification, and protein-protein binding affinity estimation. We demonstrate PROBE’s usage through a new use case evaluating ESM2 and three recent multimodal PLMs—ESM3, ProstT5, and SaProt—highlighting their ability to integrate diverse data types, including sequence and structural information. This study underscores the potential of protein language models in advancing protein function prediction and serves as a valuable tool for both PLM developers and users.

## 1. Introduction

Proteins are fundamental to life, serving as the driving force behind countless biological processes, from cellular regulation to metabolic pathways. Understanding protein characteristics is vital not only for advancing biological and evolutionary research but also for their practical applications in biotechnology and therapeutic development [1]. Annotating protein functions through experimental methods is resource-intensive, both in terms of time and cost and lags behind the rapid accumulation of sequence data generated by high-throughput technologies.

This challenge has led to the development of computational protein function prediction methods, utilising a variety of algorithms and architectures as essential tools for achieving greater scalability and accuracy in protein annotation [2–5]. One of the most widely used frameworks for protein function annotation is the Gene Ontology (GO) [6], a structured vocabulary that organises protein functions into categories based on molecular activity, biological roles, and subcellular localisation. The standardised format of GO ensures consistency in annotation across species and databases, enabling comprehensive comparative studies and more accurate predictions. The Critical Assessment of Functional Annotation (CAFA), a widely adopted benchmarking framework, has played a pivotal role in advancing the field, providing valuable insights into the strengths and limitations of current prediction techniques [7–9].

Despite these advancements, challenges remain, with ongoing efforts focusing on enhancing the accuracy of predictions related to enzymatic activities, structural properties, and the dynamics of molecular interactions [10–13].

Proteins should be quantitatively/numerically represented to be utilised in computational function prediction models. Early protein representation methods were model-driven, relying on predefined rules and homology-based techniques, such as sequence alignment and evolutionary relationships, to infer and express protein characteristics [14, 15]. These methods employed known quantitative measurements of physical, chemical, or biological properties to generate feature vectors [16, 17]. While effective for smaller datasets, their reliance on predefined rules and limited flexibility made it difficult to scale with the growing complexity of high-throughput data [7]. In response, the field has shifted towards data-driven techniques, where statistical and machine learning algorithms learn complex, high-dimensional representations directly from protein features such as sequences and structures. Both supervised and unsupervised learning algorithms are employed to train models on large-scale biomedical datasets, creating adaptive representation vectors that capture both local and global patterns. Small-scale protein representation learning methods [18–22], inspired by natural language processing techniques like word2vec [23], focused on local patterns but were limited in capturing broader contexts. Deep learning techniques, including convolutional neural networks (CNNs) [24], long short-term memory (LSTM) networks [24–28], and larger-scale transformer-based [29] architectures [24, 30–33], have since advanced to capture both local and global features, recognising long-range dependencies across entire protein sequences.

Since transformer-based models were originally developed for and used in the natural language processing field (e.g., large language models), protein representation models that utilise the same sequence-based data processing approach and machine learning architecture are generally named protein language models (PLM). Furthermore, recent multimodal representation learning techniques [34–36], integrating diverse data types like sequence and structural information, provide a more holistic view of protein characteristics, enhancing predictive accuracy. This progress has made protein representation models more flexible, scalable, and efficient for large-scale protein annotation and analysis.

Despite significant progress in protein representation methods and prior benchmarking efforts [24, 31, 37], there remain gaps in evaluating how well these models capture the multifaceted functional features of proteins. This need for comprehensive, multitask evaluation led to the development of our “Protein Representation Benchmark”–PROBE system [1]. Our study addresses this gap by evaluating a wide range of protein representation methods—both classical [14–17, 38, 39] and machine learning-based [18–21, 24–26, 30–36, 40, 41]—across four distinct benchmark tasks, each targeting a different aspect of protein function. Figure 1 provides a schematic overview of the PROBE benchmarking framework. Out of these four benchmarking tasks, “semantic similarity inference” evaluates the models’ ability to capture functional relationships between protein pairs through GO annotations. A related task, “ontology-based protein function prediction”, measures how accurately models assign specific GO terms to input proteins. The third one, “drug target family classification”, tests the models’ ability to categorise proteins into their generic superfamilies, reflecting evolutionary and functional patterns. Finally, “protein-protein binding affinity estimation” assesses how well models predict interaction changes due to mutations, offering insights into protein interaction dynamics.

**Figure 1.**
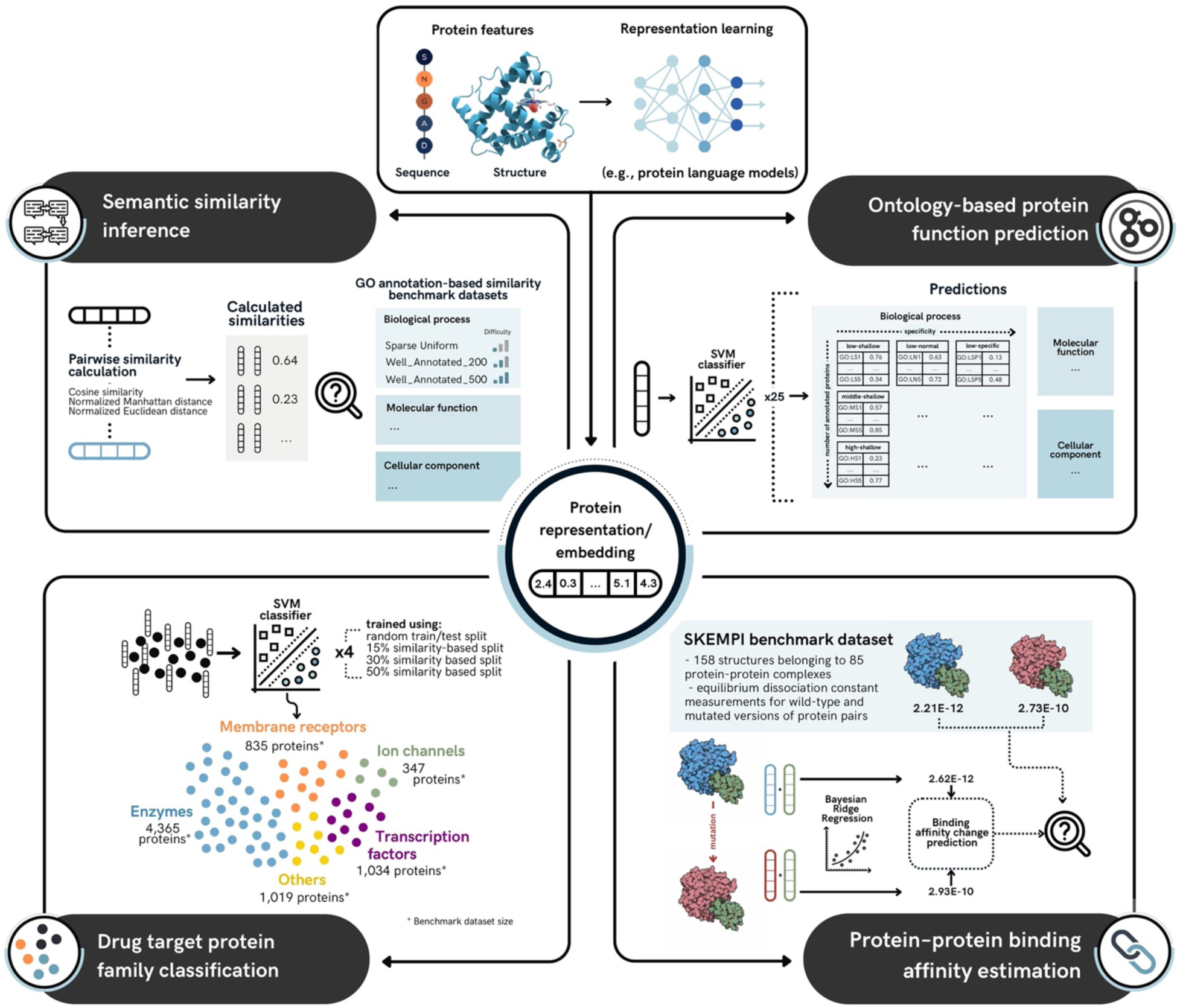
Protein representation/embedding learning process and the four benchmark tasks used to evaluate protein representation methods. Protein features such as sequence and structure are transformed into numerical vectors (embeddings) using representation models (e.g., PLMs). These embeddings are then evaluated across four benchmark tasks: **Semantic similarity inference**, which compares pairwise protein similarities to ground-truth GO-based similarities; **Ontology-based protein function prediction**, where classifiers predict protein functions across GO categories; **Drug target protein family classification**, which categorises proteins into superfamilies; and **Protein–protein binding affinity estimation**, which predicts binding affinity changes due to mutations and compares them against the ground-truth values (in the SKEMPI dataset).

We have developed the PROBE tool (https://github.com/kansil/PROBE) to enable users to assess any representation method across the four benchmark tasks and compare their results with those reported in this study. To further enhance accessibility, we introduced a user-friendly online graphical interface (https://huggingface.co/spaces/HUBioDataLab/PROBE), which is now available, accompanied by detailed usage instructions included in this paper. Additionally, we have now expanded our discussion by incorporating new results from advanced PLMs, including ESM-2 [33], ESM-3 [36], ProstT5 [34] and SaProt [35], the last three of which represent cutting-edge multimodal PLMs, reflecting the latest advancements in protein representation learning. Our benchmarking framework offers researchers a valuable resource for assessing representation models in predictive modelling and ultimately highlighting innovative approaches to help address unresolved challenges in protein science.

## 2. Materials and Methods

In our previous study [1], we conducted a comprehensive evaluation of protein representation methods to assess how effectively they capture meaningful protein features and apply this information in downstream tasks to predict functional properties. The evaluation now includes 25 methods, ranging from classical approaches (e.g., BLAST [14], HMMER [15], PFAM [38]) to recent machine learning-based models (e.g., ProtVec [18], ProtBERT [31], ESM [30]), selected for their proven success in previous predictive tasks and availability as open-access tools or pre-constructed vectors. We designed four function-related benchmarking tasks: (i) semantic similarity inference, (ii) ontology-based protein function prediction (PFP), (iii) drug target protein family classification, and (iv) protein-protein binding affinity estimation, each of which measures how well the representation methods captured features relevant to the specific problem. In the semantic similarity task, our models predicted pairwise protein relationships based on Gene Ontology (GO) annotations. For the ontology-based protein function prediction task, they predicted specific GO terms associations of proteins. In the drug target protein family classification task, models classify proteins into target superfamilies based on patterns that correspond to the defining characteristics of each family. Finally, in the protein-protein binding affinity task, models estimated changes in binding affinity values between protein isoforms/proteoforms due to mutations.

### 2.1 Datasets

#### Semantic similarity inference dataset

We downloaded human protein entries from the UniProtKB/Swiss-Prot database [42] along with their GO term annotations from the UniProt-GOA database 2019_11 release [43], retaining only human expert-reviewed annotations by excluding electronically inferred annotations (IEA).

These annotations were propagated to parent terms within the GO hierarchy, following the true path rule. The final dataset included 14,625 distinct GO terms across three categories: molecular function (MF), biological process (BP), and cellular component (CC), with a total of 326,009 annotations. Ground truth pairwise GO-based semantic similarities between proteins were calculated separately for each category using the Lin similarity measure [44] from the GoSemSim package [45], involving 3,077 proteins for MF, 6,154 for BP, and 4,531 for CC. To address potential bias from poorly annotated proteins, we created three subsets— Well_Annotated_500, Well_Annotated_200, and Sparse Uniform—for each GO category, yielding nine datasets. The Well_Annotated_500 subset focused on the top 500 proteins with the highest number of GO annotations, while Well_Annotated_200 included the top 200.

However, these subsets exhibited dense similarity score regions, where proteins with similar numbers of annotations showed very close similarity values. This dense clustering led to rank changes among pairs with proximal similarities, which significantly decreased correlation values and reduced the discriminative power of the measurements. The Sparse Uniform dataset was generated to solve this problem by sampling every thousandth protein pair from the Well_Annotated_500 set, resulting in 247 similarity scores between 40 proteins, making it easier to predict, while Well_Annotated_500 remains the most challenging due to its annotation complexity.

#### Ontology-based protein function prediction dataset

We retrieved human protein entries and their GO term annotations from the UniProtKB/Swiss- Prot and UniProt-GOA databases (release 2019_10), excluding electronically inferred annotations (IEA) to enhance the reliability of the data. For each GO term, individual protein lists were created for cross-validation and filtered using UniRef50 clusters [46] to ensure that train/test datasets excluded protein sequences with more than 50% similarity, avoiding bias in the analysis. GO terms were then grouped based on the number of annotated proteins into three categories: low (2–30 proteins), middle (100–500 proteins), and high (over 1,000 proteins). Additionally, GO terms were classified by their specificity as shallow, normal, or specific, depending on their depth within the GO hierarchy. These classifications produced nine GO term groups per GO category (MF, BP, and CC), resulting in a total of 27 groups. Two groups (MF-high-specific and CC-high-specific) were not analysed due to a lack of corresponding GO terms. For the remaining 25 groups, five GO terms were selected based on maximised dissimilarity, measured using Lin similarity, ensuring that the evaluation covered a wide range of terms with high functional diversity. Groups with fewer than five terms included all available terms.

#### Drug target protein family classification dataset

The dataset was constructed using curated drug/compound–target protein interaction data from the ChEMBL (v25) chemistry database [47]. Human drug target proteins were categorised into five groups based on ChEMBL’s hierarchical classification: four main families—enzymes, membrane receptors, transcription factors, and ion channels—and a fifth group labelled as "others" for the remaining targets. Additional human proteins were gathered from UniProt’s curated keyword and Enzyme Commission (EC) number annotations. For transcription factors, we used a comprehensive study cataloguing human transcription factors [48] and manually filtered the list to retain high-confidence members. After eliminating ambiguous or redundant family annotations, the dataset was finalised with 4,365 enzymes, 835 GPCRs, 347 ion channels, 1,034 transcription factors, and 1,019 proteins in the "others" category. The data was split into training and test sets using UniClust’s [49] pre-calculated clustering scheme at 50% and 30% sequence similarity thresholds. Additionally, following the UniClust protocol, we used the MMSeq tool [50] to generate clusters at a 15% similarity threshold. At each similarity threshold, no sequence pairs exceeded the specified similarity level between the train and test sets. To ensure fair comparisons, the test/validation sets were kept identical across splits, and sequences exceeding the similarity threshold were excluded from the training set for each fold. Family annotations were used as class labels during training.

#### Protein-protein binding affinity estimation dataset

For this benchmark task, we utilised the structural database of kinetics and energetics of mutant protein interactions (SKEMPI) dataset [51], a collection of experimentally measured data on mutation-induced binding affinity changes for protein-protein heterodimeric complexes gathered from the literature. SKEMPI provides 3,047 measurements of the equilibrium dissociation constant (KD) across 158 protein structures representing 85 different protein-protein complexes (PDB models). Each data point includes two KD values for the same protein pair: one for the wild-type version and another for a variant containing one or more single amino acid mutations. The binding affinity changes were calculated by taking the difference between KD values. To evaluate the predictive power of the protein representation methods, we used 2,950 data points from the SKEMPI dataset following the PIPR study [52], assessing how well the models could predict the binding affinity for both wild-type and mutated proteins independently.

### 2.2 Protein representation vectors

Our benchmark study covers a wide range of protein representation methods (see Note 5), from small-scale PLMs including Learned-Vec [19], Mut2Vec [20], Gene2Vec [21], TCGA_EMBEDDING [40], CPCProt [41], and ProtVec [18]; to large PLMs including SeqVec [25], TAPE-BERT-PFAM [24], MSA-Transformer [32], ProtBERT-BFD [31], UniRep [26], ESM-1b [30], ProtALBERT [31], ProtXLNet [31], and ProtT5-XL [31], which have greater than 10 million learnable parameters. In this study, we extend the set of large PLMs by adding ESM-2 (esm2_t33_650M_UR50D) [33] alongside ESM-3 (ESM3_OPEN_SMALL) [36], ProstT5 (the only available version) [34], and SaProt (SaProt35M_AF2) [35]—three multimodal approaches that integrate sequence, structure and additional information to further improve predictive performance. Table 1 summarises the protein representation learning methods, outlining their machine learning algorithms, input data types, embedding size (number of dimensions), applications, relevance, and available data repositories.

**Table 1.**
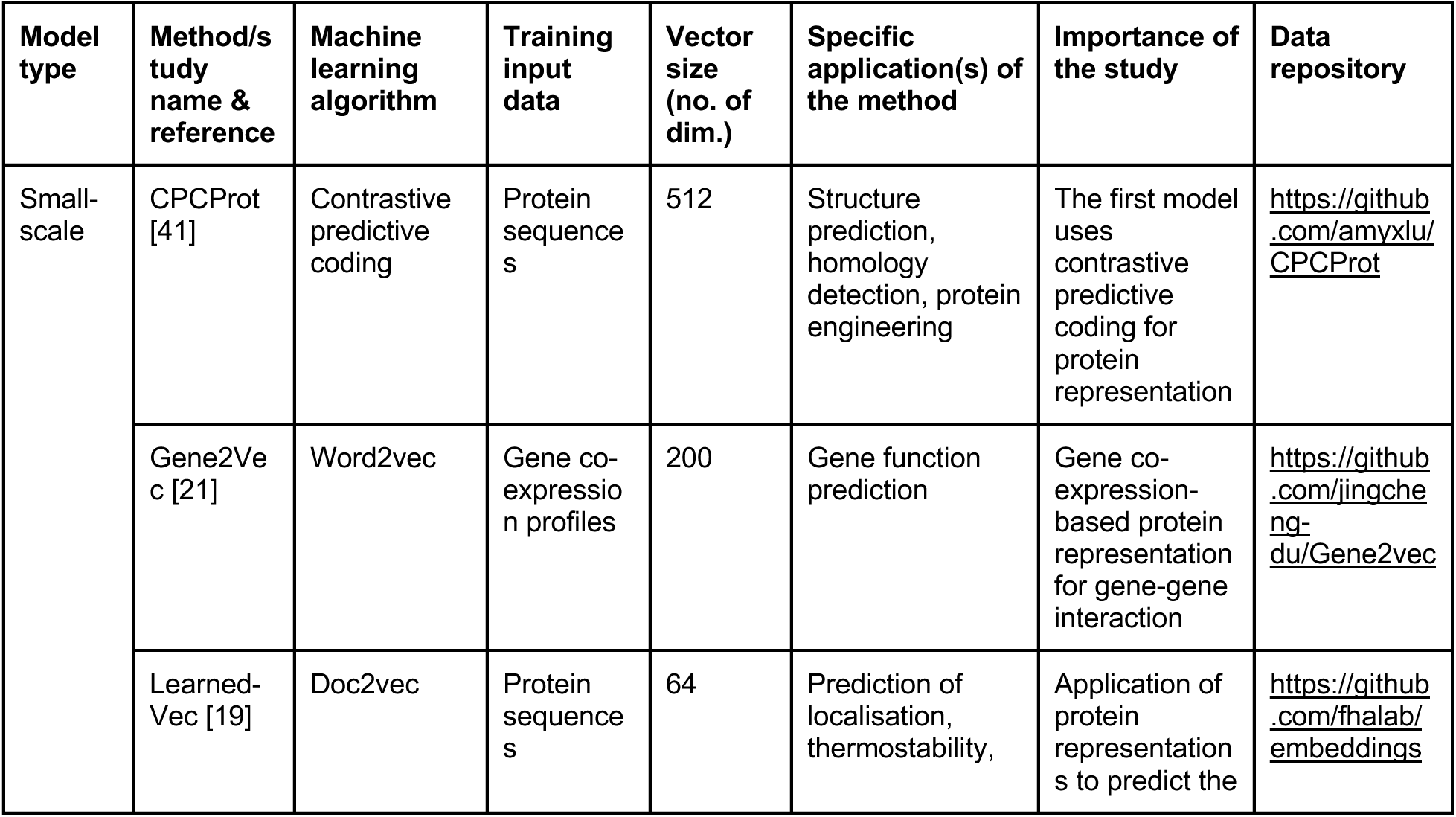
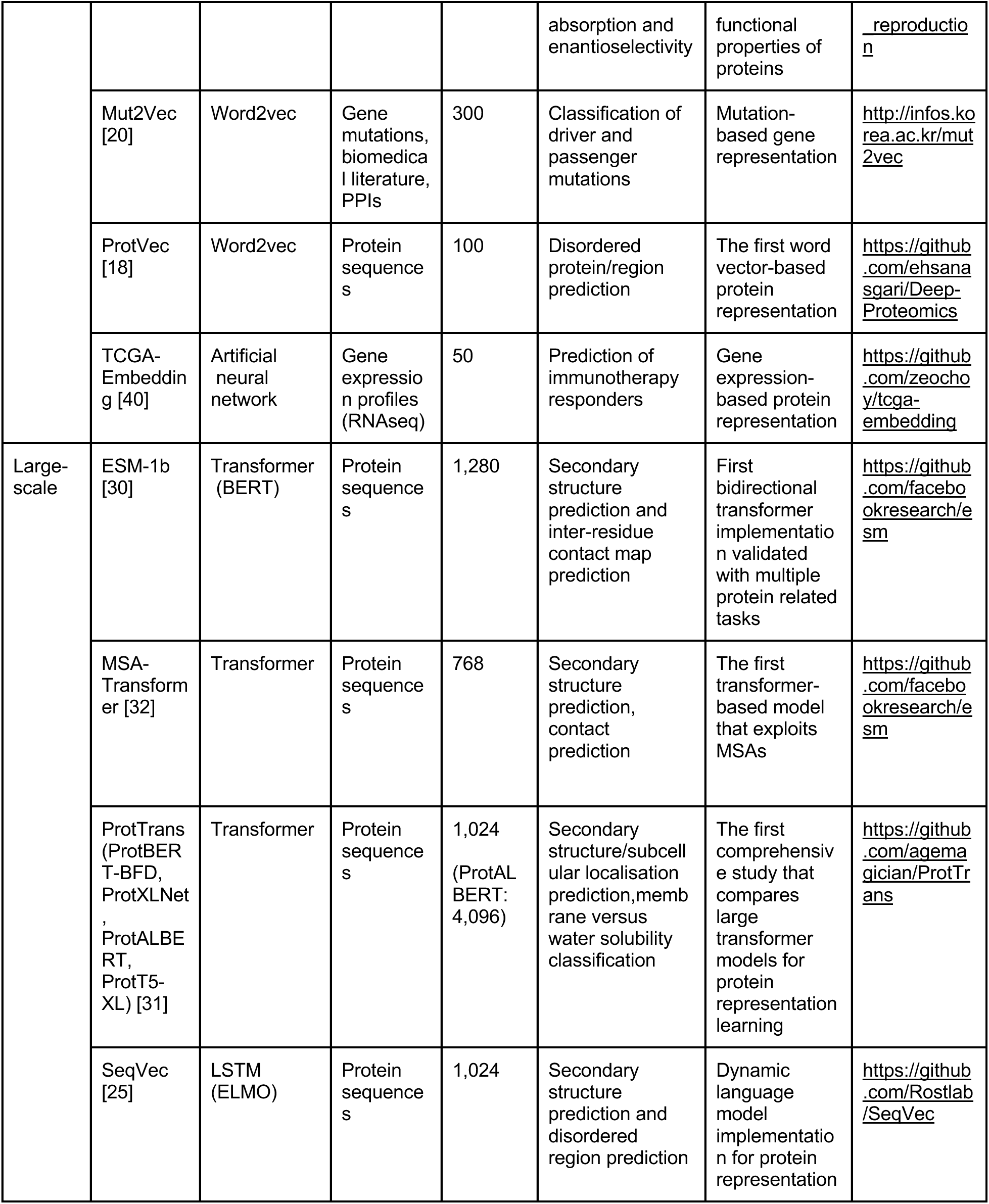
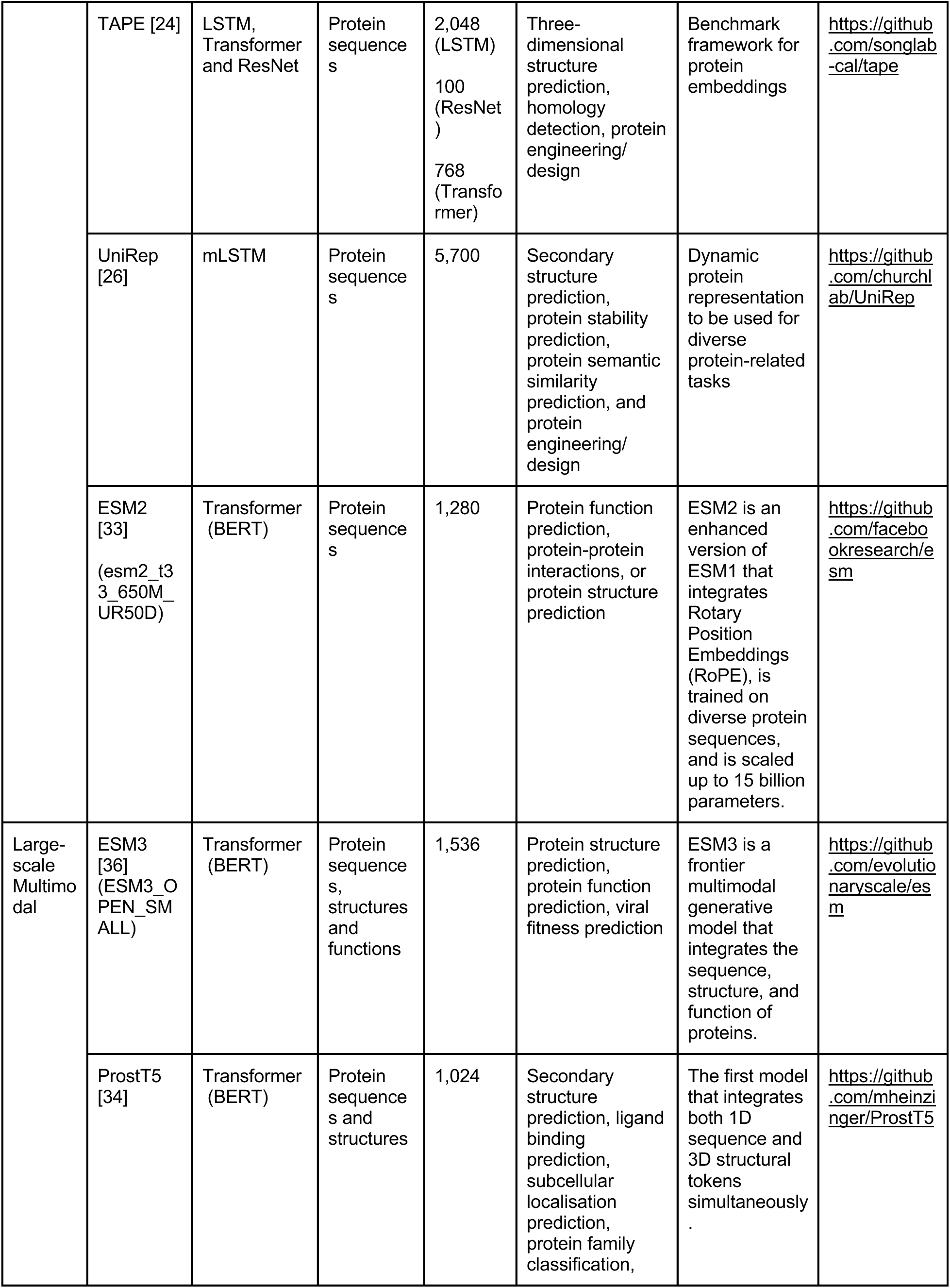
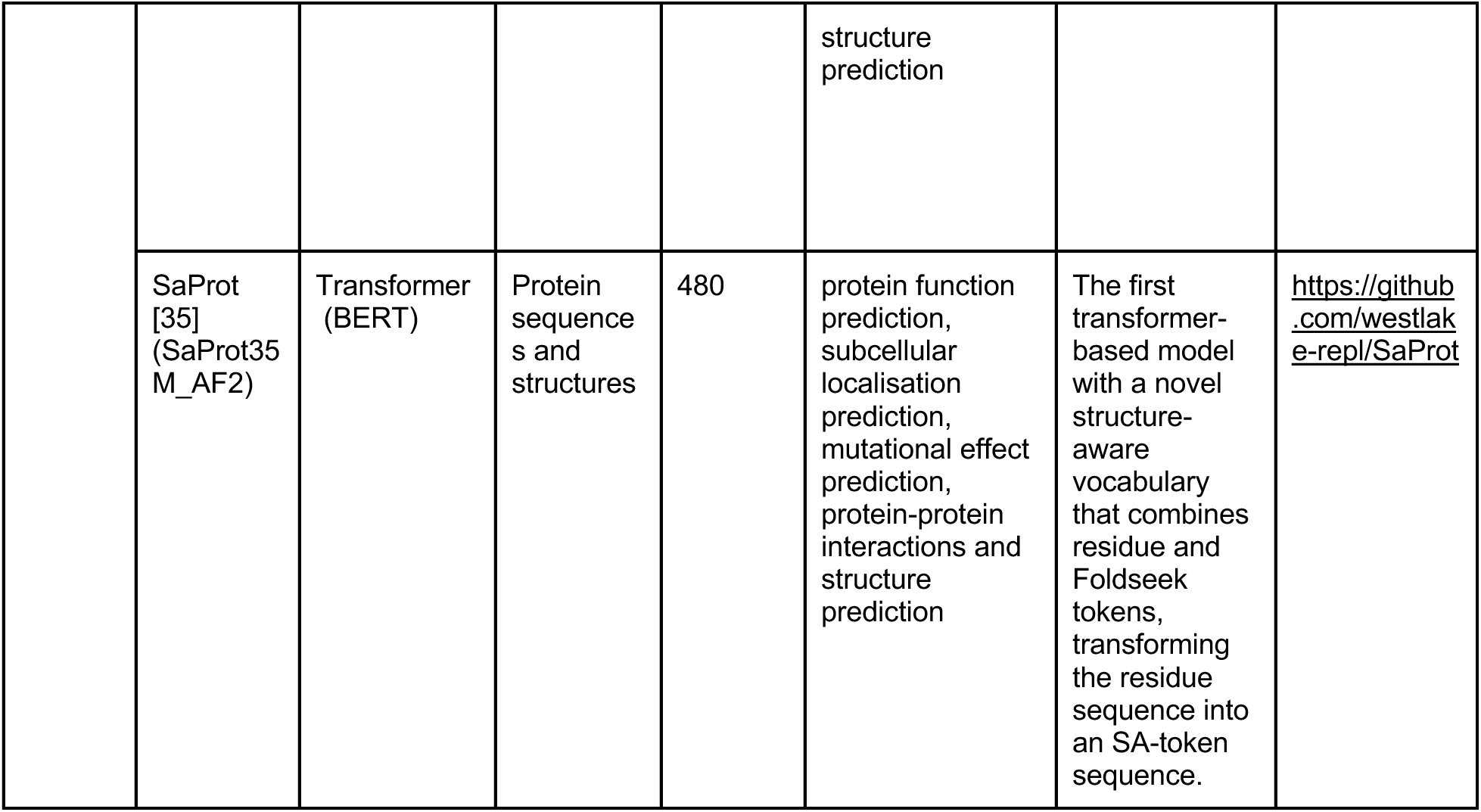
The list and details of protein representation learning methods (mostly PLMs) included in our analysis.

Classical methods such as BLAST [14], HMMER [15], AAC [39], APAAC [16], K-Sep [17], and PFAM [38] served as baselines, relying on traditional homology searches, amino acid composition, and evolutionary data. For all protein entries in our datasets, representation vectors were either sourced directly from pre-existing vector data or generated using the source code of the respective models when necessary. For the protein-protein binding affinity estimation benchmark, amino acid sequences corresponding to the complex structures in the SKEMPI dataset were retrieved from PDB, and representation vectors were computed using these sequences (see Note 4).

It is also important to mention that, in this study, we offer both a programmatic tool and an online benchmarking platform (explained in section 2.4) where the user can upload any representation vector set and run the benchmark tasks. Therefore, the framework is not limited to the models mentioned above.

### 2.3 Benchmarking Framework

For the semantic similarity inference task, pairwise similarities between the representation vectors of proteins from each method were calculated using cosine similarity, normalised Manhattan distance, and normalised Euclidean distance (the latter two converted to similarity by subtracting from 1). These were compared to ground truth pairwise semantic similarities derived from Gene Ontology (GO) annotations. Model performances were evaluated by computing Spearman rank-order correlations between vector-based similarities and true semantic similarity rankings.

For the ontology-based PFP task, we trained 500 independent linear support vector machine (SVM) classification models using stochastic gradient descent (SGD) and Hinge loss, covering 25 GO benchmark groups across 20 methods, all implemented via the scikit-learn library [53]. Each model’s performance was assessed using fivefold cross-validation with default hyperparameters.

In the drug target protein family classification task, four scikit-learn linear SVM classifiers with stochastic gradient descent (SGD) and default parameters were trained for each representation method—one for the random split and three for the similarity-based splits. The OneVsRest Classifier mode handled multiple classes, and models were trained and tested using tenfold cross-validation.

For the protein-protein binding affinity estimation task, element-wise multiplication was applied to the representation vector pairs to generate input vectors for the prediction model. To maintain consistency with prior studies, we adopted the same estimator, cross-validation strategy, and random states used in the PIPR study [52]. Bayesian Ridge Regression, implemented via scikit-learn, was used as the binding free energy estimator with tenfold cross-validation. Results were compared against other methods; Siamese residual RCNN, Siamese residual GRU, Siamese CNN, and baseline approaches such as autocovariance and composition-transition-distribution. Performance was measured using mean squared error (MSE) and mean absolute error (MAE), with ground truth values derived from the SKEMPI dataset (*see* Note 7).

### 2.4 Web service

To streamline the benchmarking process, we have developed a user-friendly web server for the PROBE platform using Gradio, an open-source interface framework (https://www.gradio.app). Our web service, accessible at https://huggingface.co/spaces/HUBioDataLab/PROBE, allows researchers to evaluate their protein representation/language models across our four distinct benchmark tasks: (i) semantic similarity inference, (ii) ontology-based protein function prediction, (iii) drug target protein family classification, and (iv) protein-protein binding affinity estimation.

The interface, designed to focus on accessibility, necessitates solely an up-to-date web browser and an internet connection. Users can upload their model’s representation vectors for the proteins in our benchmark datasets and specify the desired task(s) and sub-task(s)/dataset(s) for evaluation. The backend scripts automatically handle the benchmarking process, and results are displayed in a table format with key metrics such as Spearman rank correlation, F1-score, accuracy, mean squared error (MSE), etc., depending on the selected task.

The web service is organised into three primary sections/tabs. The main “PROBE Leaderboard” section/tab (Figure 2a) displays the results of methods in a table format, with options for users to visualise performance metrics and compare selected models through intuitive graphical representations (Figure 2b). This visual analysis aids in understanding how models perform across different benchmarks and simplifies model evaluation and refinement.

**Figure 2.**
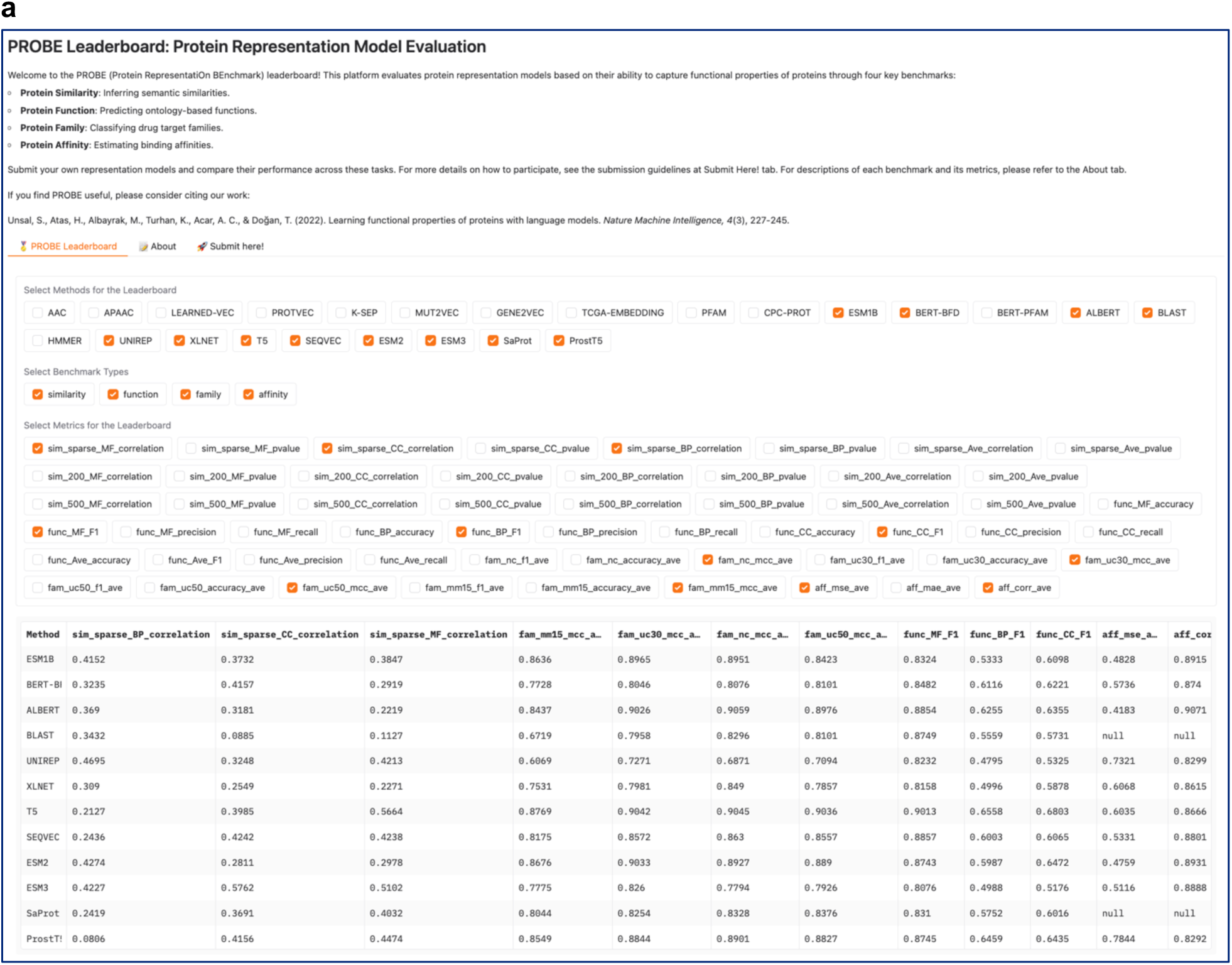
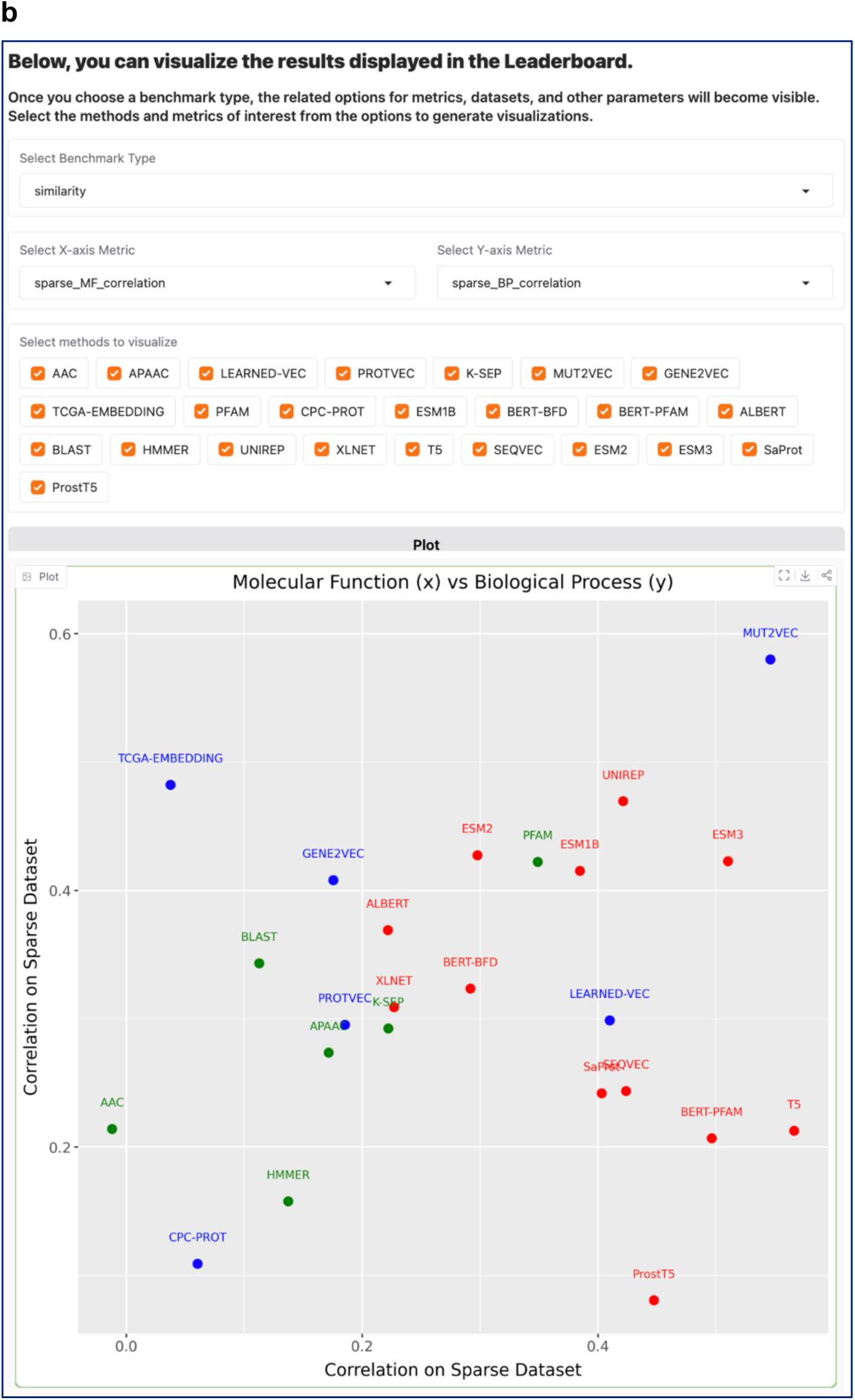
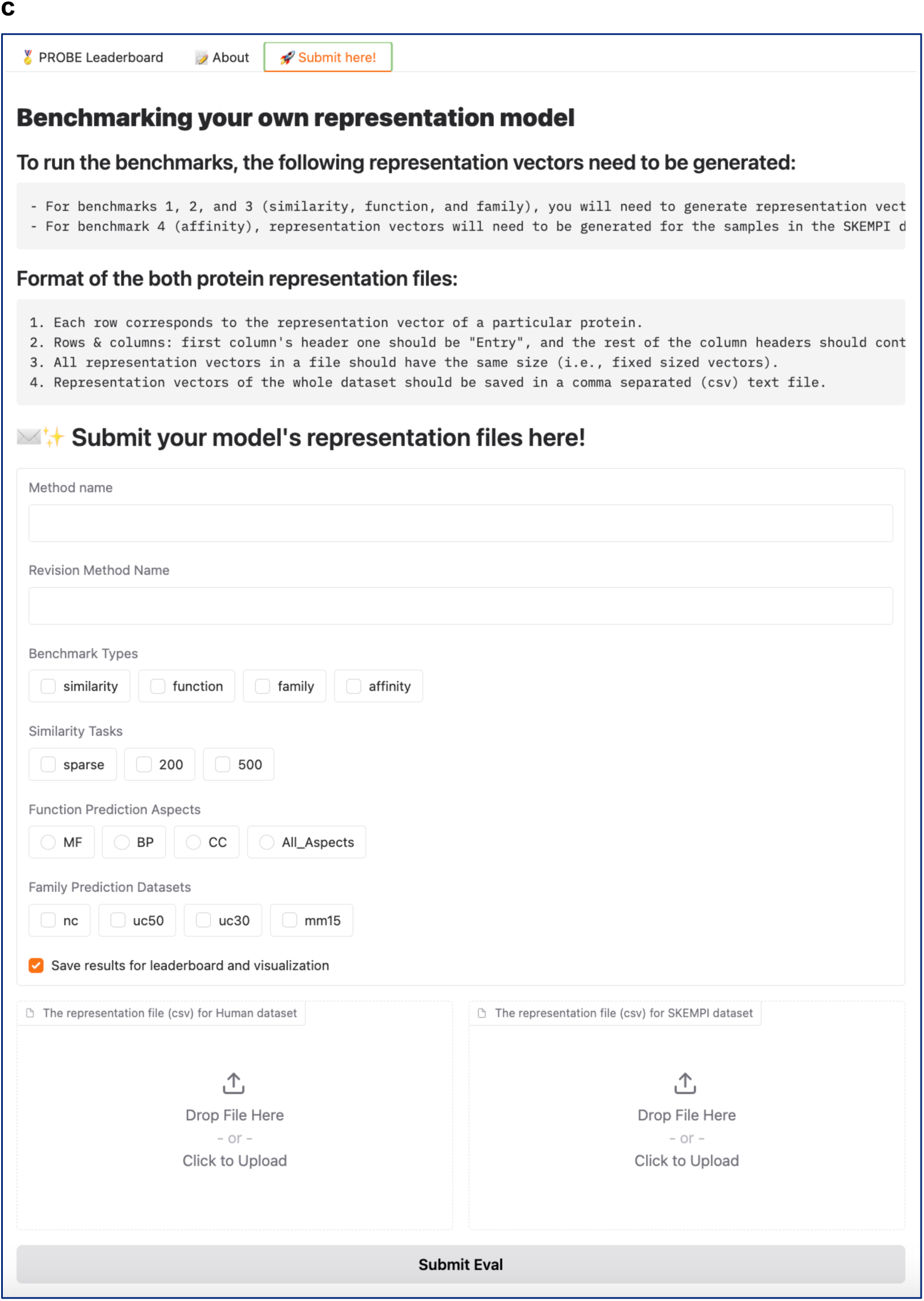
PROBE web service interface (https://huggingface.co/spaces/HUBioDataLab/PROBE). **(a)** The PROBE Leaderboard tab provides an interface for users to visualise and compare the performance of various methods on selected metrics. Users can interactively choose methods and metrics to display customised leaderboards and **(b)** generate comparative plots; **(c)** the submission tab allows users to upload new model representation files for evaluation. It offers options to select benchmark types and evaluation tasks, streamlining the submission process.

The “About” section/tab provides detailed information regarding the PROBE methodology, offering insights into the four benchmark tasks and the evaluation framework. This section serves as a resource for users seeking a deeper understanding of the principles behind the benchmarks.

The “Submit Here” section/tab (Figure 2c) is dedicated to user submissions, where researchers can upload their representation files, select specific benchmarks and datasets, and initiate the evaluation process. Once submitted, the system processes the data and presents both numerical results and visual comparisons, allowing users to assess their model’s performance across various biological tasks.

By automating complex evaluation processes, the PROBE web service ensures a seamless experience from submission to result interpretation, offering valuable insights into how well a representation model (e.g., a PLM) captures various functional aspects of proteins.

## 3. Use Case Study

In this use case, we ran the whole PROBE benchmark using the programmatic tool (https://github.com/kansil/PROBE) on four new PLMs, namely ESM2, ESM3, ProstT5, and SaProt (detailed in the last four rows of Table 1) and analysed the findings by comparing the results against protein representation methods provided in the original PROBE study (the rest of Table 1). It is important to note that the PROBE benchmark mainly aims to facilitate protein function-related tasks (see Note 1). Below, we provide detailed information about each step, together with the visualisation and discussion of results.

### 3.1 Generating representation vectors (embeddings)

We compiled protein representation vectors for the human protein entries in our dataset using the selected PLMs (i.e., ESM2, ESM3, ProstT5, and SaProt). Residue-level embedding vectors, derived from the pre-trained representation models, are aggregated using mean pooling to generate a single fixed-size representation for each protein. In the embedding file, the rows correspond to protein UniProt IDs, while the columns represent the dimensions of vectors (see Note 3). The final embedding/representation vector files can be accessed at https://drive.google.com/drive/folders/1Mx8CHs2yJQCKpC02fIX_ZxCrN1H8ZQ64. We obtained the pre-constructed representation vectors of other models using the links provided in the GitHub repo.

### 3.2 Installation, configuration and running of the benchmark

To run the PROBE benchmark via the GitHub repo (https://github.com/kansil/PROBE), we first cloned the repository and accessed the project directory. We installed any necessary dependencies and downloaded the required datasets and/or representation vectors. The downloaded files were placed in the specified directories or paths, as outlined in the PROBE repository documentation (https://github.com/kansil/PROBE/blob/master/README.md). The configuration file (Figure 3) was adjusted before executing the benchmark (https://github.com/kansil/PROBE/blob/master/bin/probe_config.yaml). This configuration file enables flexible customisation, allowing users to efficiently switch between models and datasets to assess different aspects of protein representation methods. The representation_name field specifies the protein representation models to be incorporated, such as ESM2 or ProstT5. The corresponding tasks are selected using the benchmark option, which allows the choice between semantic similarity inference (similarity), ontology-based protein function prediction (function), drug target protein family classification (family), and protein-protein binding affinity estimation (affinity). Four benchmarks are simultaneously run by selecting all. The paths to the protein embedding files are defined in the representation_file_human and representation_file_affinity fields, where the respective CSV files containing the embedding vectors are provided. For semantic similarity inference, the specific datasets to be used can be selected via similarity_tasks. In the case of function prediction, the Gene Ontology (GO) aspect is chosen using function_prediction_aspect, while the dataset size splits are selected through function_prediction_dataset. Similarly, for drug target protein family classification, similarity-based splits are defined using family_prediction_dataset. To generate comprehensive results, the detailed_output option can be set to True. Following the configuration procedure, we ran all benchmarks and datasets on selected representation models (i.e., ESM2, ESM3, ProstT5, and SaProt, together with models from the original study for comparison purposes) (see Note 2 for runtimes).

**Figure 3.**
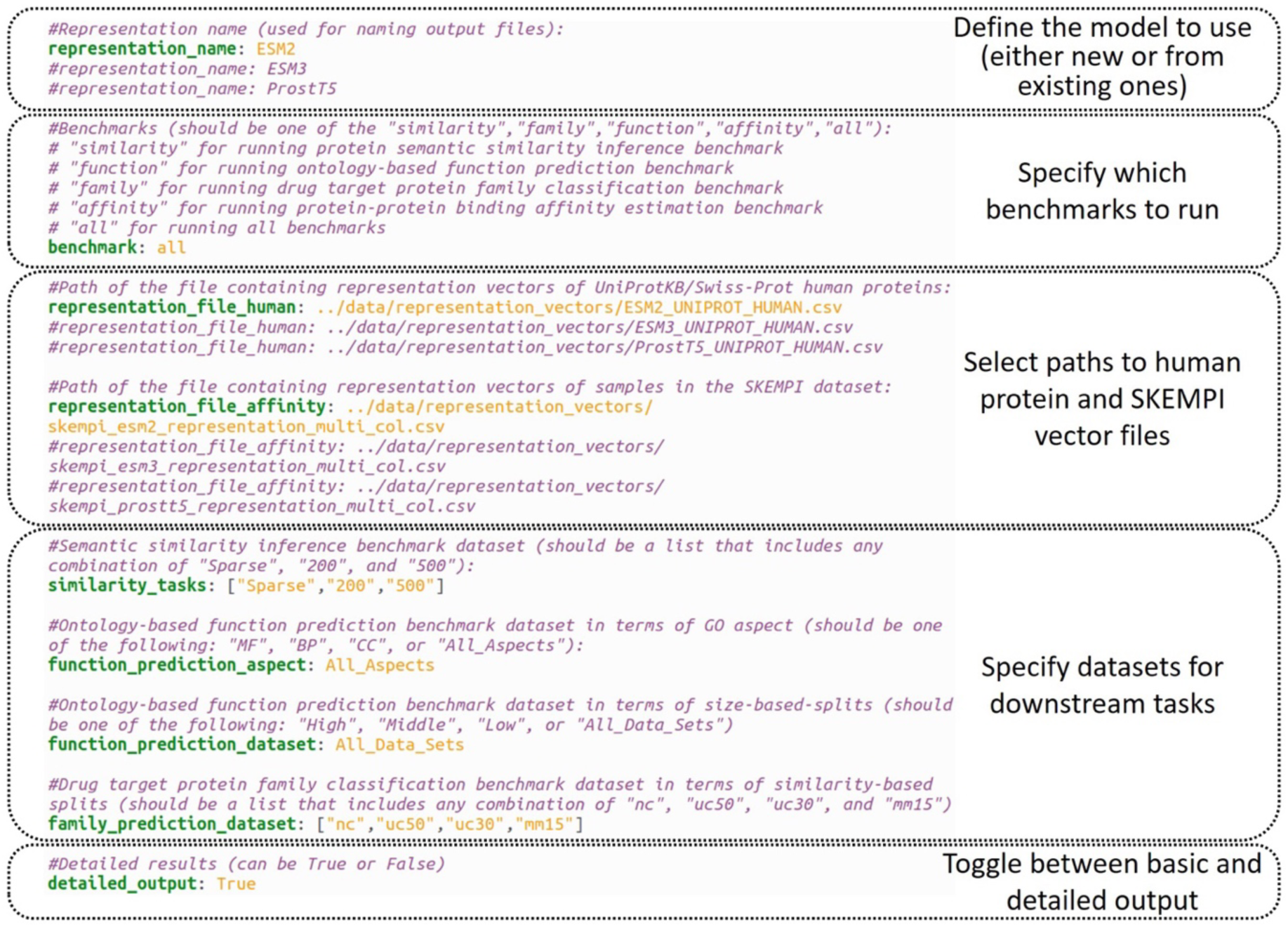
An example configuration file. The figure illustrates the structure and components necessary for running the benchmark.

### 3.3 Analysing the results

#### Semantic similarity inference

This analysis evaluates the extent to which representation models capture biomolecular functional similarity. Spearman rank-order correlation values were calculated to assess performance by comparing the ranked lists of representation vector similarities with the true GO-based semantic similarities. Results based on the Manhattan distance are given in Figure 4.

**Figure 4.**
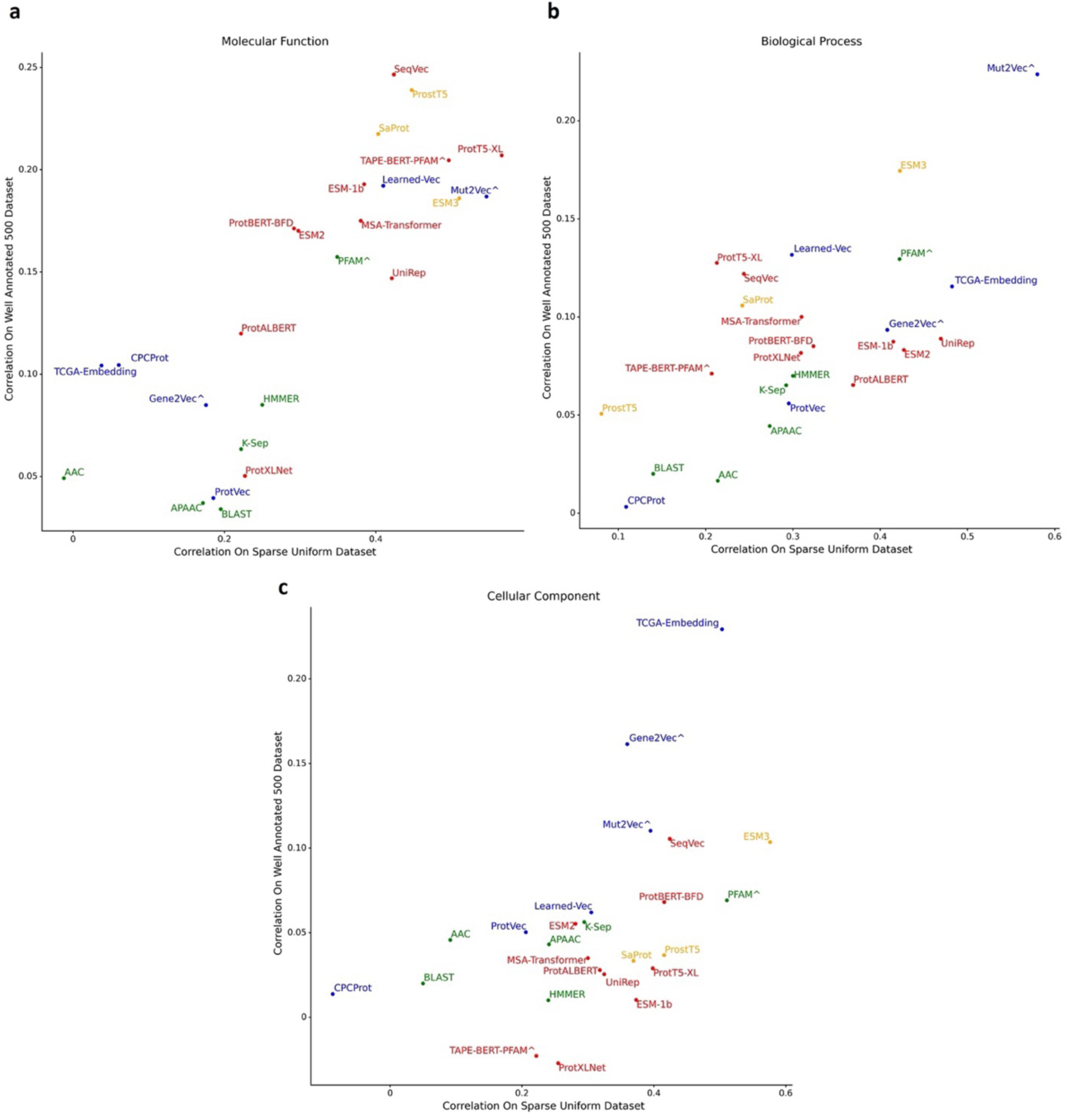
Protein semantic similarity inference benchmark results. Performance (Spearman correlation) of protein representation methods in inferring pairwise semantic similarities between proteins in Spearman correlation calculated on Manhattan distances, considering GO categories of **(a)** molecular function, **(b)** biological process, and **(c)** cellular component, on Sparse Uniform (x-axis) and Well_Annotated_500 (y-axis) datasets (model names; green: classical representations, blue: small-scale PLMs, red: large-scale PLMs, and orange: multimodal PLMs). Methods with data leak suspicion are marked by ^ symbols.

Considering the GO MF category, ProtT5-XL demonstrated the highest performance within the Sparse Uniform dataset, while SeqVec outperformed the remaining models on the well-annotated 500 set, with ProstT5 and SaProt following closely. Mut2Vec consistently achieved the best results across both datasets in the GO BP category. However, there is a data leak suspicion for this model, which highlights the performances of ESM3 and TCGA-Embedding. In the GO CC category, TCGA_EMBEDDING and ESM3 achieved the highest correlation scores.

To summarise, ESM3 excelled in capturing semantic similarities across all GO categories. SaProt and ProstT5 were also good at GO MF. These models leverage multimodality, allowing them to simultaneously capture sequence and structure-related features and identify complex relationships within protein data, which may have contributed to their strong performance.

Incorporating Foldseek [54] structural tokens probably yielded a critical structural insight.

#### Ontology-based protein function prediction

Gene Ontology (GO) term annotations were utilised to benchmark protein representation models using an SVM classifier. The overall prediction performance, as illustrated in the F1- score heat maps in Figure 5, suggests a superior predictive outcome for the GO MF category compared to GO BP and CC for most models. ProstT5, ESM2, ESM3, and SaProt model performances are similar to each other across all GO categories (MF, BP, and CC), delivering top-tier results in both the Biological Process (BP) and Cellular Component (CC) categories. ProtT5-XL ranked highest in MF performance while performing well in other GO categories.

**Figure 5.**
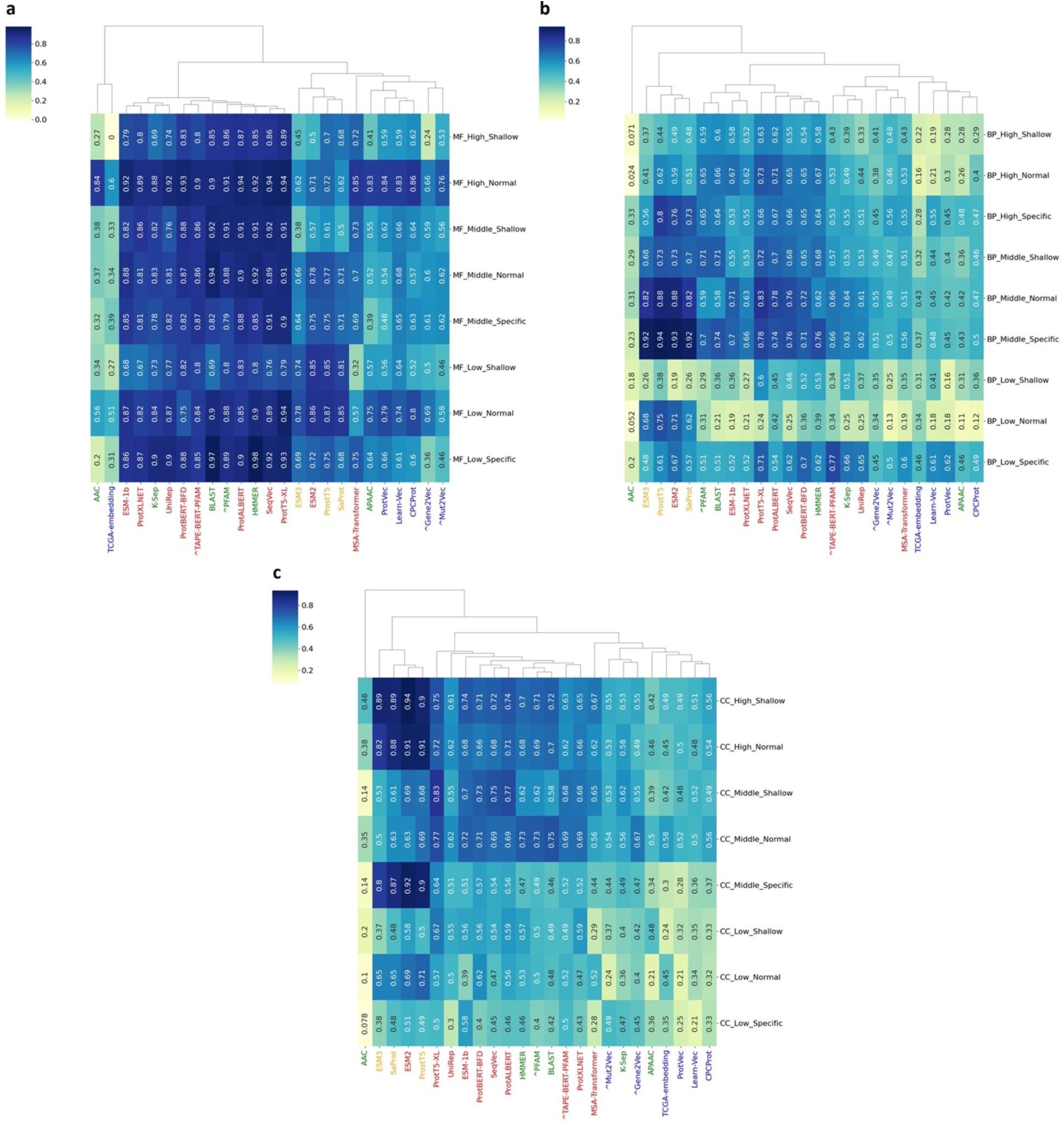
Ontology-based protein function prediction benchmark results. Heat maps indicating the clustered performance results in terms of weighted F1-scores of protein representation methods in ontology-based PFP benchmark for GO categories of **(a)** molecular function, **(b)** biological process, and **(c)** cellular component (model names; green: classical representations, blue: small-scale PLMs, red: large-scale PLMs, and orange: multimodal PLMs). Methods with data leak suspicion are marked by ^ symbols.

ProstT5 excelled in BP, and ESM2 achieved the best results in CC. Notably, ProtT5-XL, ESM2, and ProstT5 share the transformer architecture but differ in design—ESM2 is encoder-only, while ProstT5 and ProtT5-XL’s encoder-decoder structure enables conditional language modelling. Similar to semantic similarity inference, Multimodal models generally performed well in this task.

#### Drug target protein family classification

We evaluated protein representation performance in drug discovery by predicting the main/superfamilies of drug target proteins, i.e., enzymes, membrane receptors, transcription factors, ion channels and others, using four different sequence similarity-based cut-offs for train-test splits. The overall performance was assessed using the F1-score, accuracy, and Matthews correlation coefficient (MCC). According to the results shown in Figure 6, ProtT5-XL is the top performer across all datasets, except for the random split, where it ranks second. Upon evaluating the performance metrics, the highest-ranked models across all datasets exhibited comparable results, regardless of whether multimodality was utilised. In other words, multimodality did not lead to an improvement in drug target protein family classification performance (see Note 6).

**Figure 6.**
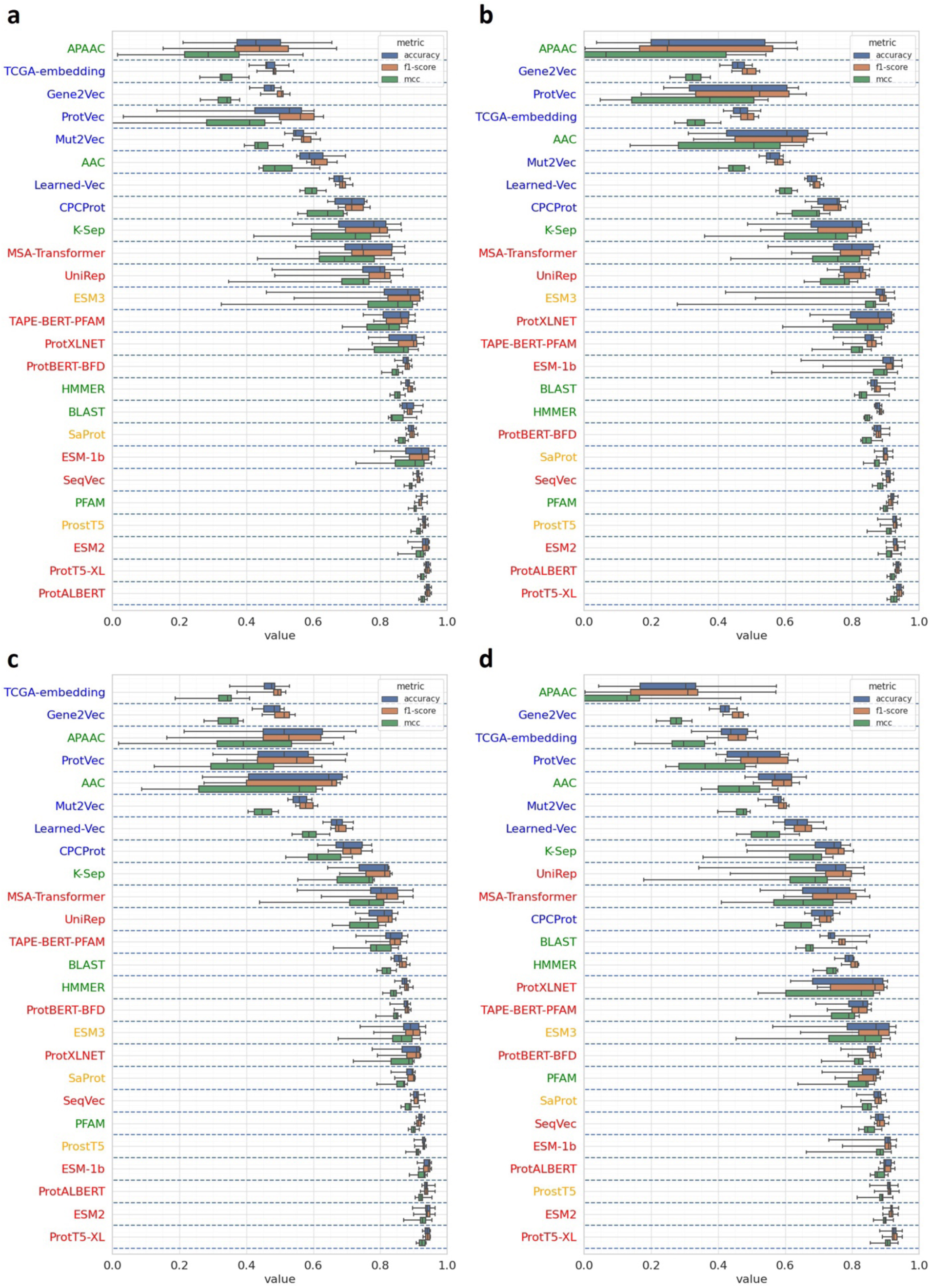
Drug target protein family classification benchmark results. Box plots displaying the overall performance results in F1-score, accuracy and MCC of protein representation methods in the drug target protein family classification benchmark on **(a)** the random split, **(b)** UniClust50, **(c)** UniClust30, and **(d)** MMSEQ-15 datasets. Models are sorted according to mean MCC scores (model names; green: classical representations, blue: small-scale PLMs, red: large-scale PLMs, and orange: multimodal PLMs). Whiskers indicate minimum/maximum values.

### Protein-protein binding affinity estimation

We evaluated protein representation methods considering their performance in estimating the protein-protein binding affinity changes between the wild-type proteins and their mutated versions by employing Pearson correlation, mean squared error (MSE), and mean absolute error (MAE) (Figure 7). Here, ProtALBERT delivered the best predictions, significantly outperforming the baseline PPI prediction model, Siamese residual RCNN (PIPR), by about 25%. Furthermore, the ESM-1b, ESM2, and ESM3 models consistently outperformed the baseline across all evaluation metrics and took close-to-top places. Notably, the high performance of nearly all transformer-based models indicated that the attention mechanism could effectively capture amino acid interactions. Here, one exception is the ProstT5 model, which displayed a comparatively poor performance, while its base model, ProtT5, scored decently. The reason might be that ProstT5 was not designed as a general-purpose PLM. As mentioned in the original paper, even though the authors took precautions, catastrophic forgetting might have occurred during training, reflected in performance drops in the prediction of binding residues, conservation, and subcellular locations [34].

**Figure 7.**
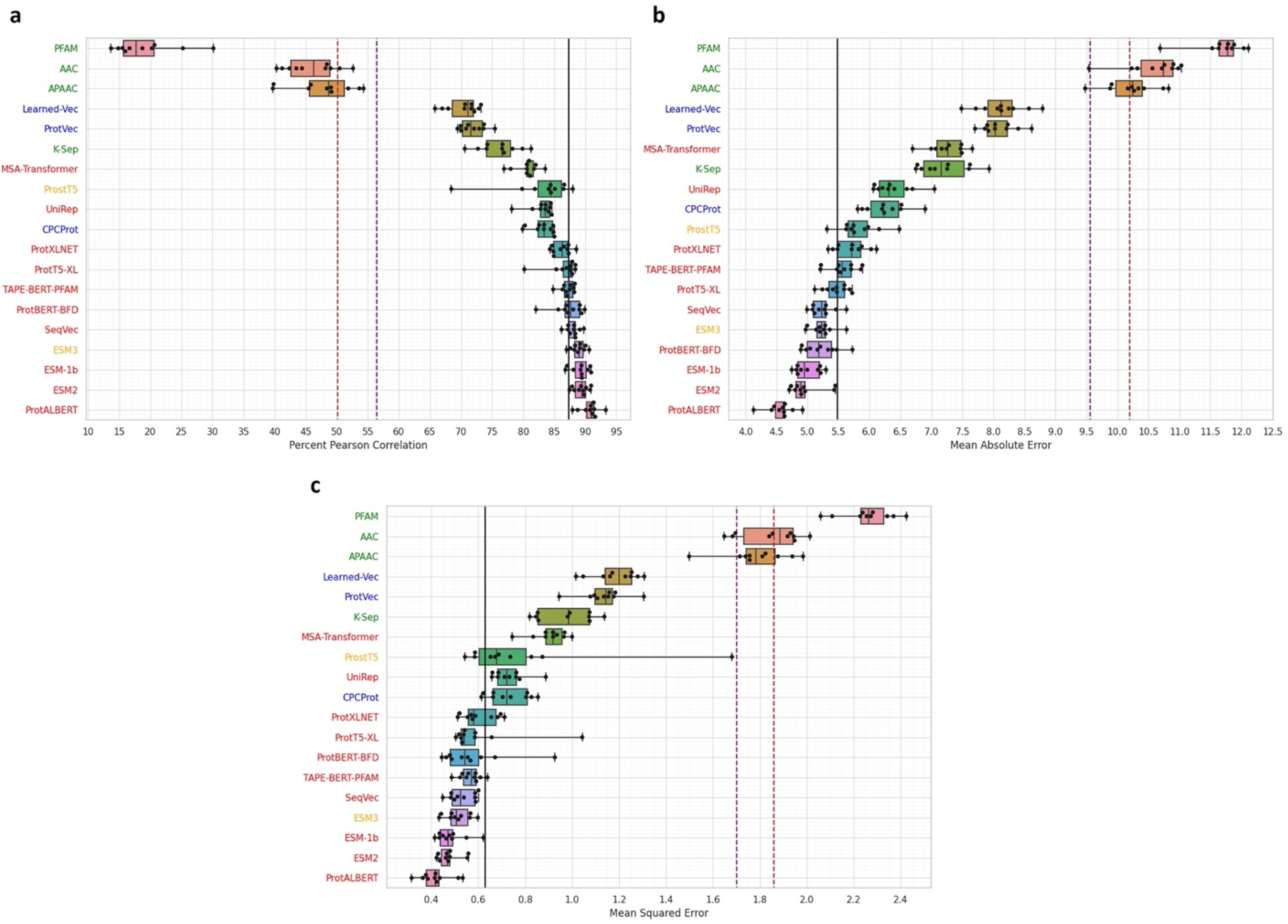
Protein-protein binding affinity estimation benchmark results. Box plots indicating performance results of protein representation methods in the protein-protein binding affinity estimation benchmark: **(a)** % Pearson correlation values indicating the correlation between predicted values and the true binding affinities where higher values are better, **(b)** MSE values where lower values are better, **(c)** MAE values where lower values are better. Each dot shows the performance of a fold in the ten-fold cross-validation (model names; green: classical representations, blue: small-scale PLMs, red: large-scale PLMs, and orange: multimodal PLMs). The black vertical line indicates the best, and the dashed lines indicate baseline scores in the PIPR [52] study. Whiskers indicate minimum/maximum values. Box colours represent specific models and provide for easy comparison between panels.

## 4. Notes

1. PROBE is a benchmarking tool that evaluates and compares different protein representations based on their performance in function-related prediction tasks. It is not designed as a generalised protein function prediction tool. For this purpose, our methods such as Domain2GO [55] (https://huggingface.co/spaces/HUBioDataLab/Domain2GO), UniGOPred [56] (https://unigopred.kansil.org/), ECPred [57] (https://ecpred.kansil.org/), and DEEPred [2] can be used.
2. We have recorded the runtimes and resource utilisation of PROBE while evaluating different PLMs. The assessment includes embeddings derived from human protein and SKEMPI datasets across all benchmarks available in PROBE, except for SaProt. For SaProt, we exclusively run benchmarks on human proteins. The experiments were conducted on our local server with the following hardware configurations: CPU (Intel Xeon Gold 5215) with 64GB DD4 RAM, on the Ubuntu 20.04 operating system. Table 2 summarises the profiling of the new PLMs utilised in our use case study. The resource utilisation, specifically CPU and memory, was systematically recorded at different points during runs. The table reports the maximum values in the recorded data.
3. The PROBE tool is designed to handle two primary input files: representation vectors for human proteins (for three of our benchmarking tasks) and the SKEMPI dataset proteins (for the binding affinity prediction benchmarking task). Users can use either or both datasets to run the benchmarks using the PLM of their choice (choosing from the representation methods included in our study). If the user wishes to test a new representation model / PLM, the vectors of the proteins in the selected dataset(s) should be generated externally and fed to our system.

**Table 2.** Maximum CPU and memory usage, run time, and embedding sizes for newly incorporated protein language models (PLMs) in the PROBE benchmark evaluation. CPU usage is calculated in terms of % of a single thread.

| PLM (number of parameters) | CPU usage (%) | Memory usage (MB) | Run time (sec) | Embedding size (dimension) |
| --- | --- | --- | --- | --- |
| ProstT5 | 94.9% | 1235.28 | 1490.87 | 1024 |
| ESM2 | 227.7% | 1431.38 | 1643.63 | 1280 |
| ESM3 | 155.8% | 1649.08 | 1905.50 | 1536 |
| SaProt | 224.7% | 588.07 | 1182.45 | 480 |

### Protein sequences utilised in our study

Human protein dataset sequences can be accessed at: https://drive.google.com/file/d/1wXF2lmj4ZTahMrl66QpYM2TvHmbcIL6b/view

SKEMPI protein sequences can be accessed at: https://drive.google.com/file/d/1m5jssC0RMsiFT_w-Ykh629Pw_An3PInI/view

It is also important to note that PROBE is capable of benchmarking any numerical protein representation; therefore, non-PLM-based representations (e.g., physicochemical descriptors) can also be used. Representation vectors should satisfy the conditions below.

### Requirements for representation/embedding vectors

- **Aggregation:** Though most PLMs generate per-residue embeddings, PROBE requires a single aggregated embedding for the entire sequence. For the aggregation method, please refer to the documentation of the chosen PLM. This is usually not an issue for non-PLM-based representations, as most work at the protein level.
- **Uniformity in Embedding Size:** The generated vectors must be uniform in size for any single method. Note that PROBE can accommodate different embedding sizes across different methods.
- **Header Information:** The first column’s header should be labelled “Entry” for the file representing the human proteins and “PDB_ID” for SKEMPI proteins. Subsequent column headers should be indices corresponding to the dimensions of the representation vector.
- **Content:** The first column below the header needs to list the IDs for the respective proteins. Each following column must contain the numerical values corresponding to the embedding features for each dimension. That is, each row corresponds to a unique protein.
- **File Type:** Representation data should be stored in a comma-separated (CSV) file.
- **Example Data:** The following link contains embedding vectors that PROBE can process: https://drive.google.com/drive/folders/1Mx8CHs2yJQCKpC02fIX_ZxCrN1H8ZQ64
4. Most existing protein representation models are trained using a single type of data, usually the protein sequences. However, protein knowledge encompasses various biological information types, such as protein-protein interactions (PPIs), post-translational modifications, gene/protein (co)expressions and so on. There is a growing trend toward models integrating multiple data types (i.e., multi-modal models), particularly those incorporating structural information. In our original benchmark, the majority of models were based solely on sequence data. In this study, we included three multi-modal models, namely ESM3, SaProt and ProstT5.
5. This benchmark is mostly centred on evaluating protein representation learning models in the context of protein functions, from Gene Ontology annotations to protein families. For tasks related to structural feature prediction, i.e., residue/angle/distance prediction and inverse folding, it is recommended to refer to alternative benchmarks such as ProteinWorkshop [58] (https://github.com/a-r-j/ProteinWorkshop).
6. The performance of top models often converges on high-value regions (i.e., > 0.9 in terms of F1 score and MCC) in tasks such as Gene Ontology (GO) based protein function prediction (considering molecular function) and drug-target protein family classification. While evaluating a new model, this situation makes it difficult to observe whether the new model offers superior predictive capabilities compared to the existing top methods. In other words, it is not possible to significantly outperform the existing top performers in the abovementioned tasks because there is not much room for further improvement. In these cases, in order to distinguish the performance difference between the best models, it would be a better choice to pay more attention to biological process and cellular component-based GO and semantic similarity prediction or protein-protein binding affinity estimation tasks.
7. Some methods had to be omitted from the protein-protein binding affinity estimation benchmark. BLAST and HMMER were excluded because this task utilises protein variants with incomplete sequences (fragments), and BLAST and HMMER feature vectors were created using the complete wild-type sequences. Mut2Vec, Gene2Vec, and TCGA_EMBEDDING, which are genomic data-based methods lacking protein-related information, were discarded. Finally, SaProt was also omitted because of the missing 3D structural data for mutated proteins.

